# Septin 7 Interacts With Numb To Preserve Sarcomere Structural Organization And Muscle Contractile Function

**DOI:** 10.1101/2023.05.11.540467

**Authors:** Rita De Gasperi, Laszlo Csernoch, Beatrix Dienes, Monika Gonczi, Jayanta K. Chakrabarty, Shahar Goeta, Abdurrahman Aslan, Carlos A. Toro, David Karasik, Lewis M. Brown, Marco Brotto, Christopher P. Cardozo

**Affiliations:** Departments of Psychiatry and Friedman Brain Institute, Icahn School of Medicine at Mount Sinai; Medicine, Icahn School of Medicine at Mount Sinai; Rehabilitation Medicine, Icahn School of Medicine at Mount Sinai; Spinal Cord Damage Research Center, James J. Peters VA Medical Center; Department of Physiology, Faculty of Medicine, University of Debrecen, Hungary; ELKH-DE Cell Physiology Research Group, University of Debrecen, Debrecen, Hungary; Quantitative Proteomics and Metabolomics Center, Department of Biological Sciences, Columbia University; Azrieli Faculty of Medicine, Bar Ilan University, Safed, Israel; Bone-Muscle Research Center, College of Nursing & Health Innovation, University of Texas at Arlington

## Abstract

Here, we investigated mechanisms by which aging-related reductions of the levels of Numb in skeletal muscle fibers contribute to loss of muscle strength and power, two critical features of sarcopenia. Numb is an adaptor protein best known for its critical roles in development including asymmetric cell division, cell-type specification and termination of intracellular signaling. Numb expression is reduced in old humans and mice. We previously showed that, in mouse skeletal muscle fibers, Numb is localized to sarcomeres where it is concentrated near triads; conditional inactivation of Numb and a closely related protein Numb-like (NumbL) in mouse myofibers caused weakness, disorganization of sarcomeres and smaller mitochondria with impaired function. Here, we found that a single knockout of Numb in myofibers causes reduction in tetanic force comparable to a double Numb, NumbL knockout. We found by proteomics analysis of protein complexes isolated from C2C12 myotubes by immunoprecipitation using antibodies against Numb, that Septin 7 is a potential Numb binding partner. Septin 7 is a member of the family of GTP-binding proteins that organize into filaments, sheets and rings, and is considered part of the cytoskeleton. Immunofluorescence evaluation revealed a partial overlap of staining for Numb and Septin 7 in myofibers. Conditional, inducible knockouts of Numb led to disorganization of Septin 7 staining in myofibers. These findings indicate that Septin 7 is a Numb binding partner and suggest that interactions between Numb and Septin 7 are critical for structural organization of the sarcomere and muscle contractile function.

## INTRODUCTION

A common consequence of aging is sarcopenia, which is characterized by declining skeletal muscle mass, muscle contractile force, muscle power and overall physical function [1]. The causes of reduced contractile function and power of skeletal muscle in individuals who have sarcopenia remain incompletely understood. One candidate mechanism is the reduced expression of the adaptor protein Numb in muscle of older organisms. Numb mRNA expression is reduced in muscle biopsy samples during normal aging in humans [2] and Numb protein levels are diminished in skeletal muscle of 24 month old mice [3]. Numb is an adaptor protein that is highly conserved from Drosophila to humans. Numb contributes to asymmetric cell division, specification of cell fate, trafficking of cell surface proteins such as integrins, and turnover of signaling molecules such as Notch, Hedgehog and p53 [4–6]. A role in mitochondrial fission and fusion has also been suggested in certain contexts [7]. In mammals, the Numb gene contains 10 exons and is expressed as one of four splice variants. The three shorter variants lack exons 3, 9 or both. Removal of exon 3 shortens the phosphotyrosine binding domain while removal of exon 9 truncates a proline-rich domain (Supplementary Figure 1).

Within skeletal muscle, Numb has dual roles. Its expression in satellite cells is critical for their proliferation and for tissue repair after muscle injury [8]. In skeletal muscle fibers, a conditional knockout (cKO) from myofibers of Numb and the closely related protein Numb-like (NumbL) reduced muscle contractility [3]. Examination of muscle from mice with a double-knockout of Numb and NumbL by transmission electron microscope revealed perturbed muscle ultrastructure and altered mitochondrial morphology [3]. Ultrastructural changes included increased spacing of Z-lines, staircasing of Z-lines, altered mitochondrial morphology and loss of the regular spacing of sarcoplasmic reticulum [3]. *In-vitro* experiments using mouse primary myotube cultures found that knockdown (KD) of Numb reduced myoblast fusion and mitochondrial function, and delayed caffeine-induced calcium release [3]. A subsequent study found that knockout (KO) of Numb in the heart led to cardiac dilation and altered cardiac and skeletal muscle sarcomere structure [9]. Since NumbL protein levels are very low or undetectable in adult skeletal muscle, we have posited that the effects of KD of Numb and NumbL on skeletal muscle ultrastructure and function are likely attributable to loss of Numb expression [3].

Molecular mechanisms by which Numb and NumbL organize the sarcomere and assure optimal calcium release during excitation-contraction coupling remain poorly understood. The interactome of Numb includes p53 [10, 11], Mdm2 [12, 13], the LNX family of proteins which target Numb for degradation by the ubiquitin-proteasome pathway [14–16], and proteins involved in internalization and trafficking of membrane proteins, Eps 15 and α-Adaptin [17]. Co-immunoprecipitation (Co-IP) experiments have shown that Numb binds sarcomeric α-actin and actinin [9] and Numb has been proposed to participate in sarcomere assembly [9]. The possibility that Numb binds other proteins present in myofibers has not been tested.

The goal of the current study was to better understand the molecular mechanisms by which the cKO of Numb in skeletal muscle myofibers perturbed muscle weakness. We began by comparing effects of a single knockout (sKO) of Numb or a double knockout (dKO) of Numb and NumbL on *ex-vivo* physiological properties of the *extensor digitorium longus* (EDL) muscle. We then examined Numb protein binding partners in C2C12 myotubes using a combination of immunoprecipitation (IP) and liquid chromatography coupled with mass spectrometry (LC/MS/MS) approaches. Our results identify Septins as Numb binding partners and provide evidence that loss of Numb perturbs the organization of Septin 7 within myofibers.

## METHODS

### Animals

All animal studies were reviewed and approved by the James J. Peters Institutional Animal Care and Use Committee, and were conducted in accordance with requirements of the PHS Policy on Humane Care and Use of Laboratory Animals, the Guide and all other applicable regulations. C57BL/6NCrl mice were obtained from Charles River Laboratories (strain # 027). Transgenic mice in which Numb and NumbL can be conditionally ablated in skeletal muscle by injection of tamoxifen were previously described and are referred to as HSA-MCM/Numb(f/f)/NumbL(f/f) [18]. To generate mice in which a conditional, inducible KO of Numb in myofibers could be achieved, these mice were backcrossed with C57BL/6NCrl mice and bred until mice heterozygous for the HSA-MCM cassette and homozygous for the floxed-Numb allele were obtained. KD of Numb/NumbL or Numb was done by intraperitoneal injection of tamoxifen 2 mg/day for 5 days with one additional injection at day 10. The animals were sacrificed 14 days after the beginning of the induction. Controls were injected with vehicle (peanut oil). All mice were genotyped using genomic DNA isolated from ear snips as described [18]

### Numb Protein Detection by Western Blot

Numb expression was evaluated by western blotting [18, 19]. *Tibialis anterior* (TA) and EDL were homogenized using an MP homogenizer in RIPA buffer (#9806, Cell Signaling Technologies Inc., Danvers, MA) supplemented with a protease and phosphatase inhibitor cocktail (Halt, Thermo Fisher Scientific). The lysates were centrifuged at 14,000rpm for 15 minutes and the supernatant saved. Protein concentration was determined with BCA reagent (Thermo Fisher Scientific). Fifty micrograms of protein were separated by SDS-PAGE and transferred to a polyvinylidene difluoride membrane using the Trans-Blot Turbo transfer pack (Bio-Rad Laboratories, Hercules, CA). Membranes were blocked for 1 h at room temperature in Tris buffer saline 1% Tween-20 (TBS-T), 5% low-fat dry milk (blocking solution) and incubated overnight at 4°C with a rabbit monoclonal anti-Numb antibody (#2756, Cell Signaling Technology, Inc., Danvers, MA) at 1:1000 dilution in 5% BSA in TBS-T. Membranes were washed in TBST and incubated with HRP - conjugated anti-rabbit IgG at 1:2000 dilution in blocking solution (#7074, Cell Signaling Technology). Bands were revealed with ECL Prime reagent (Cytiva, Lifesciences, Piscataway, NJ) and imaged using the Amersham ImageQuant 800 imaging system (Cytiva). The blots were stripped and incubated with a rabbit monoclonal anti-GAPDH (1:1000 dilution, Cell Signaling #5174) as loading control. Western blots were quantified using ImageQuant TL software (Cytiva)

### Tissue Harvest and Ex-vivo Physiology

Animals were weighed then anesthetized using inhaled 3% isoflurane. Hindlimb muscles were excised after careful blunt dissection. Measurement of whole-muscle contractile and mechanical properties was performed using an Aurora Scientific *ex-vivo* physiology system for mice (Aurora, Ontario, Canada). A 4-0 silk suture was tied to the proximal and distal tendons of intact right EDL, immediately distal to the aponeuroses, the muscles were dissected and immediately placed in a bath containing a Krebs mammalian Ringer solution at pH 7.4, supplemented with tubocurarine chloride (0.03 mM) and glucose (11 mM) for 10 min. The bath was maintained at 25°C and bubbled constantly with a mixture of O2 (95%) and CO2 (5%). The distal tendon of the muscle was then tied to a dual-mode servomotor/force transducer (Aurora Scientific, Aurora, Ontario, Canada) and the proximal tendon tied to a fixed hook. Using wave pulses delivered from platinum electrodes connected to a high-power bi-phasic current stimulator (Aurora Scientific, Aurora, Ontario, Canada) each EDL was stimulated to contract. The 610A Dynamic Muscle Control v5.5 software (Aurora Scientific, Aurora, Ontario, Canada) was used to control pulse properties and servomotor activity, and record data from the force transducer. Optimal length (Lo) was established for each EDL to develop an isometric twitch force. EDL muscles were stimulated with a single electrical pulse to produce a twitch response. Stimulation produced a maximal twitch response by adjusting small increments (or decrements) to longer (or shorter) lengths. Muscle was left resting at least 45 s between twitch responses. Lo was achieved when twitch force was maximal. A frequency-force relationship was established once Lo was achieved. Here, EDL muscles were stimulated at increasing frequencies (i.e. 10, 25, 40, 60, 80, 100, and 150 Hz). Stimulation was delivered for 300 msec, and muscles were left to rest for 1 min between successive stimuli. Maximum absolute isometric tetanic force (Po) was determined from the plateau of the frequency-force relationship. Muscle fatigue resistance during repetitive stimulation at 60Hz every second for a total of 100 stimuli (fatigue index; FI) was also evaluated. Muscles were then removed from the bath solution and weighed. All data collected were analyzed using the Dynamic Muscle Analysis v5.3 software (Aurora Scientific, Aurora, Ontario, Canada).

### Cell Culture

C2C12 cells were grown in DMEM supplemented with 10% fetal bovine serum (FBS) and antibiotics (Penicillin-Streptomycin (10,000 U/ml, ThermoFisher) until confluency was reached then switched to DMEM supplemented with 2% horse serum and antibiotics to induce differentiation and formation of myotubes. Cells were differentiated for five days, washed three times in phosphate-buffered saline solution (PBS) and harvested with a cell scraper. At harvesting time, the cells had fused to form myotubes. The pellets were kept at -80 °C until used.

### Numb Co-Immunoprecipitation

Cell pellets were lysed in 25 mM Tris HCl pH 7.2, 150 mM NaCl, 1 mM EDTA, 5% glycerol, 0.1% Triton X-100 supplemented with Halt protease and phosphatase inhibitor cocktail (ThermoFisher) (IP buffer) for 30 minutes at 4 °C with occasional mixing. The lysate was centrifuged at 14,000 rpm for 20 minutes and the supernatant saved. Protein concentration was determined with the BCA reagent as per the manufacturer instructions (ThermoFisher).

The extracts (1 mg protein) were pre-cleared with control 4%-Agarose resin (ThermoFisher) for 1h at 4 °C with constant rotation. The samples were centrifuged and the supernatant was immunoprecipitated with 1 μg of a goat polyclonal anti-Numb (Abcam, Ab4147) which has been previously used in IP [20, 21] or with 1 μg of goat IgG (R&D Systems, Ab108-e) as IP specificity control for 1.5 hrs at 4 °C under constant rotation. To capture the complexes, 25 μl of Protein A/G Plus (Santa Cruz, sc-2003) were added and the samples incubated as above for 75 minutes. The samples were centrifuged and the beads washed twice with IP buffer, 3 times with 25 mM Tris HCl, pH 7.2, 150 mM NaCl (TBS) and once with 4 mM HEPES in 10 mM HCl. Elution of the immunoprecipitated proteins was performed by heating the samples at 37 °C for 1h with 0.3% SDS. The samples were centrifuged and supernatants were stored at -80°C.

Seven independent C2C12 samples were immunoprecipitated for the MS analysis, each pair (control IgG and anti-Numb) prepared on a different day from different cell culture samples. The Numb immunoprecipitated material was analyzed by SDS-PAGE and gels stained with silver stain kit (Bio-Rad)

### Verification of Numb Immunoprecipitation

Small scale immunoprecipitations were performed in parallel with the main IP to verify Numb IP. Proteins were separated by SDS-PAGE and blotted onto PVDF membranes. The membranes were blocked for 1 hr at 4 °C in 50 mM Tris HCl buffer, 0.15 M NaCl, 1% Tween-20 (TBS) supplemented with 0.5% non-fat dry milk (blocking solution), and incubated overnight at 4 °C with a rabbit monoclonal anti-Numb antibody (Cell Signaling #2756) diluted 1:1000 in TBS/5% BSA. After washing with TBS, the blot was incubated with HRP-conjugated anti-rabbit IgG (1:8,000 dilution in blocking solution, Cytiva), the blot developed with ECL Prime reagent (Cytiva) and imaged with an Amersham ImageQuant 800 imager (Cytiva).

### Preparation of Immunoprecipitated Samples for LC/MS/MS Analysis

The volume of the immunoprecipitated samples was adjusted to 400 μl with MS grade water and 400 μl of methanol and 100 μl of chloroform were added. The samples were mixed with a vortex mixer for 1 min and centrifuged at 14,000 rpm for 1 min. The upper phase was removed without disturbing the proteins at the interphase and 400 μl of methanol were added. The samples were centrifuged at 14,000 rpm for 5 minutes to pellet the proteins. The pellet was washed 3 times with ice-cold methanol, dried for 10 min at room temperature, resuspended in 30 μl of freshly made 100 mM ammonium bicarbonate, 8 M urea, 0.1M DTT and flash frozen in liquid nitrogen until analyzed. Cysteines were reduced and alkylated, and samples were loaded onto an S-trap micro column (Profiti C02-micro-80) according to the manufacturer’s recommendations. Proteins were digested in the trap with trypsin, eluted and lyophilized.

### LC/MS/MS Analysis

The analysis was performed using Q Exactive HF (Orbitrap) mass spectrometer coupled to an UltiMate 3000 ultra-high-performance liquid chromatography (ThermoFisher). The peptides were separated with reversed-phase chromatography using a 75 μm ID x 50 cm Acclaim PepMap reversed phase C18, 2 µm particle size column and eluted from the Nano column with a multi-step acetonitrile/formic acid gradient. The flow rate was Flow rate was 300 nL/min.

For each sample, a 2.7 h LC/MS/MS chromatogram was recorded in data-dependent acquisition mode. The 15 precursor ions with the most intense signal in a full MS scan were consecutively isolated and fragmented to acquire their corresponding MS/MS scans. The Full MS and MS/MS scans were performed at resolution of 120,000 and 15,000 respectively. S-Lens RF was set at 55%, while the Nano-ESI source voltage was set at 2.2kV.

The data were analyzed with MaxQuant_1.6.17.0. Peptide and fragments masses were searched against a database using the Andromeda search engine and scored using a probability-based approach that included a target decoy false discovery rate (FDR). Data were then analyzed with Perseus 1.6.10.50. Searches were conducted against the UniProtKB Release 2019_07 (31-Jul- 2019) (*Mus musculus* sequences, reviewed database with isoforms: 25,316 sequences) and included horse serum sequences, trypsin, keratins and common lab contaminants.

### Septin-7 Co-Immunoprecipitation

To obtain independent confirmation of Numb association with Septin 7, Numb IP was performed with anti-Numb antibody as described above. For this analysis we used three independently obtained samples of C2C12 myotubes different from those used for LC/MS/MS. The immunoprecipitated material was analyzed by Western blot using anti-rabbit polyclonal anti-Septin 7 (Proteintech, 13818-1-AP, 1:1000 dilution). To confirm Numb IP the blot was then probed with anti-Numb antibody (Cell Signaling #2756). Secondary detection was performed with Clean-Blot IP detection reagent (ThermoFisher, 21230, 1:1500).

### Analysis of Splice Variants

At least four major Numb variant forms resulting from the alternative splicing of exon 3 and/or 9 have been found [22]. To analyze Numb gene splice variants in muscle, total RNA was isolated from control and denervated gastrocnemius and from both undifferentiated and differentiated C2C12 using the Trizol reagent (ThermoFisher), further purified by the RNeasy kit (Qiagen) and reverse transcribed with the High Capacity cDNA reverse transcription reagents (Life Technologies). The cDNA was amplified by PCR using primers 5’TTCCCCCGTGTCTTTGACAG and 5’GTACCTCGGCCACGTAGAAG that span exon 1-6 to analyze exon 3 splicing and primers 5’ CTTGTGTTCCCAGATCACCAG and 5’ CCGCACACTCTTTGACACTTC spanning exon 8-10 [23] to analyze exon 9 splicing. PCR was performed using 1 μl of cDNA and Top Taq DNA polymerase and buffer (Qiagen). The reactions were performed for 30 s at 94 °C, 45 s at 57 °C and 50 s at 72 °C for 30 cycles. Aliquots of the PCR products were analyzed by 2% agarose gel electrophoresis. PCR products were cloned using the TOPO TA cloning system (ThermoFisher) and multiple resulting clones were sequenced to confirm that the expected products were generated.

### Immunohistochemical Staining

TA muscle from C57Bl/6 mice was snap-frozen in isopentane pre-cooled in liquid nitrogen. Longitudinal sections were cut a cryostat. Sections were fixed for 7 minutes in cold 4% paraformaldehyde in PBS, washed 5 times with PBS and blocked for 1 h at room temperature in TBST/0.3% Triton X-100, 5% normal goat serum (blocking buffer). The sections were then incubated overnight with rabbit polyclonal anti-Numb (1:150 dilution, Cell Signaling #2756) and with a rat monoclonal anti-Septin 7 (1:100 dilution, clone 19A4, MABT 1557, Millipore Sigma, Burlington, MA) in blocking buffer. Sections were washed with PBS and incubated with Alexa488- conjugated anti rabbit IgG (A11008, ThermoFisher) and Alexa568-conjugated anti rat-IgG (A11077, ThermoFisher) both at 1:300 dilution in blocking buffer for 2 h at room temperature. The slides were washed in PBS, stained with DAPI (1 mg/ml in PBS) and mounted with Fluorogel mounting medium (EMS, Hatfield, PA), Immunostaining was visualized by with a Zeiss LSM980 confocal microscope.

### Localization of Septin 7 in Neuromuscular Junctions

Enzymatically isolated single muscle fibers were fixed with 4% PFA for 20 minutes at RT. After the fixation 0.1 M glycine in PBS was used to neutralize excess formaldehyde. Fibers were permeabilized with 0.5% Triton-X in PBS (PBST) for 10 minutes, blocked with a serum-free Protein blocking solution (DAKO, Los Altos, CA, USA) for 30 minutes and rinsed three times with PBST solution. Anti-Septin7 (JP18991, IBL, Hamburg, Germany) diluted in blocking solution was added and the fibers incubated overnight at 4 °C in a humid chamber. Samples were washed three times with PBST and incubated with Alex-Fluor 488 conjugated alpha-bungarotoxin (1:500 dilution, B13422, Thermo Fisher) and Cy-3 conjugated anti rabbit IgG at (dilution 1:300) (A10520, Thermo Fisher) for 1 hour at room temperature. After washing three times drop slides were mounted with DAPI containing mounting medium (H-1200-10, Vector Laboratories, Burlingame, CA, USA). Images were acquired with a Zeiss AiryScan 880 laser scanning confocal microscope.

### Effect of Numb/NumbL cKO on Septin 7 Protein Expression

Hindlimb muscles from HSA-MCM/Numb(f/f)/NumbL(f/f) mice were dispersed by digestion with collagenase 1 as previously described [24]. Fibers were isolated at day 14 after inducing Numb/NumbL KD with tamoxifen as described above. Individual fibers were briefly fixed as described previously [18], incubated with anti-Septin 7 and anti-Numb antibodies as above and visualized by Zeiss LSM 700 confocal microscope.

### GWAS and WGS Searches

We conducted systematic searches of GWAS and WGS data (GWAS Catalog https://www.ebi.ac.uk/gwas/home; and Musculoskeletal Knowledge Portal, MSK-KP at https://msk.hugeamp.org/ [25]) for links between genetic variants in each of the genes encoding proteins identified as putative Numb-interacting proteins and diseases/phenotypes of interest.

### Statistics

Data are expressed as mean value ± standard deviation (STD). The significance of differences between groups was determined using either ANOVA or unpaired, 2-tailed t-tests as described in the figure legends. Statistical calculations were performed with Graphpad Prism. A p-value of < 0.5 was used as the cutoff for significance.

## RESULTS

### Effect of Numb and Numb/NumbL cKO on Force Generation

To begin, we compared properties of EDL muscle during *ex-vivo* testing between mice with a single cKO of Numb or a double cKO of Numb and NumbL. As expected, Numb protein levels in muscle lysates were significantly lower in tamoxifen-treated HSA-MCM/Numb(^f/f)^) mice and HSA-MCM/Numb^(f/f^/NumbL^(f/f)^) mice at fourteen days after starting tamoxifen, as compared to vehicle-treated mice of the same genotype (Supplementary Figure 2). Next, *ex-vivo* contractile properties of the EDL muscle were compared for HSA-MCM/Numb(^f/f)^) mice and HSA-MCM/Numb^(f/f^/NumbL^(f/f)^ mice at 14 days after starting induction with tamoxifen. A single cKO of Numb (HSA-MCM/Numb(^f/f)^) mice) markedly reduced tetanic force and twitch force generation (Figure 1A, B). There was a genotype effect of single Numb cKO for fatigue index (Figure 2 A) while time to peak tension and half relaxation time were not changed (Figure 2 B, C). As expected, EDL muscle from mice with a double Numb-NumbL cKO (HSA-MCM/Numb^(f/f^/NumbL^(f/f)^ mice) demonstrated reductions in tetanic specific tension and maximum twitch force (Figure 1 A, B) without any change in time to peak tension, half-relaxation time or fatigue index (Figure 2 A-C). There was no apparent difference for either specific tetanic tension or maximum twitch force when comparing single HSA-MCM/Numb(^f/f)^) and double (HSA-MCM/Numb^(f/f^/NumbL^(f/f)^ mice) cKO mice. These data indicate that most if not all of the reduction of force generating capacity observed in the Numb/NumbL double cKO line is attributable to inactivation of the Numb gene.

**Figure 1.**
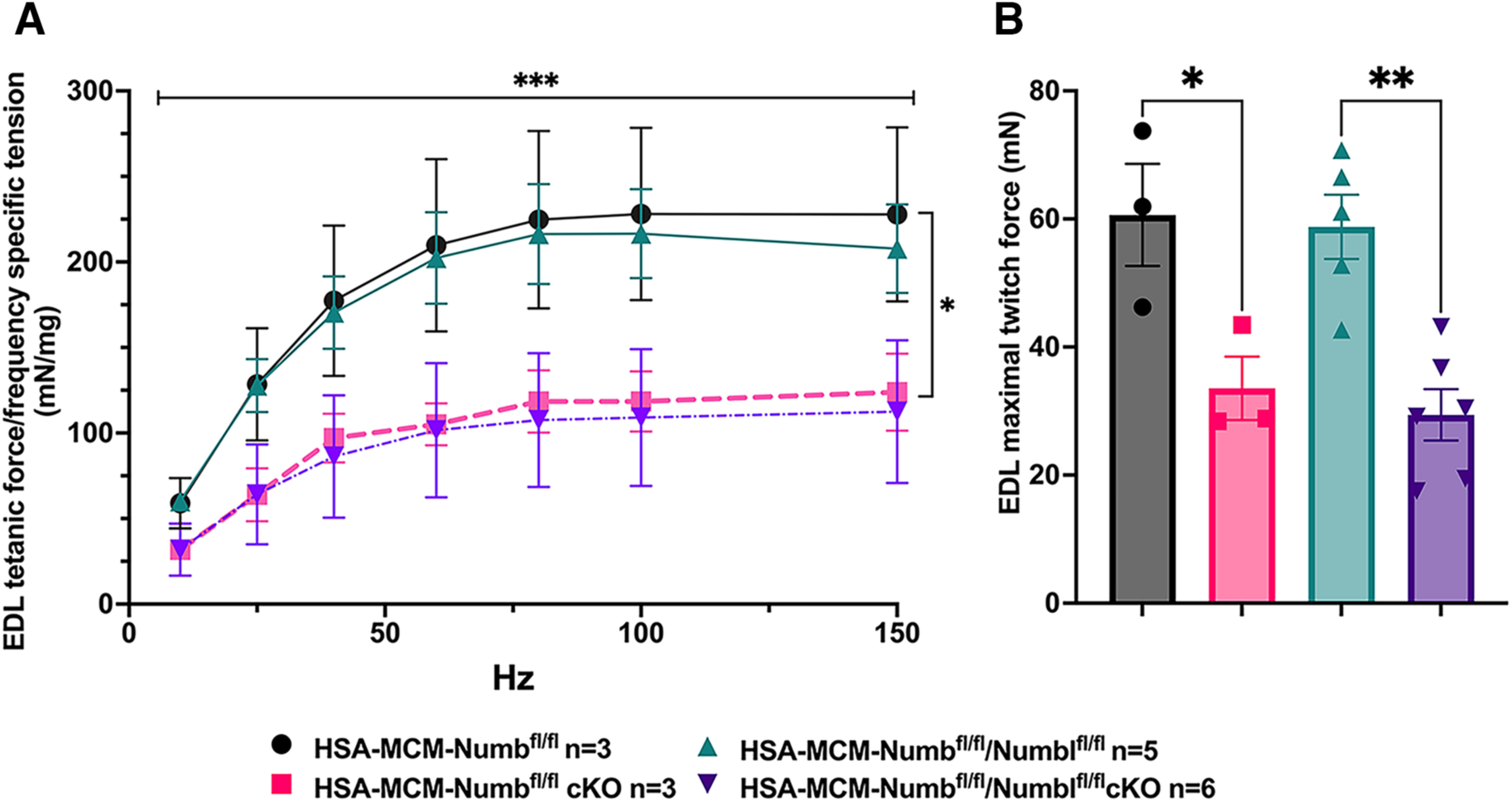
Effect of Numb or Numb/NumbL cKO on contractile function of EDL muscle during ex-vivo physiological testing. (A) Specific tension generated during tetanic contraction at the indicated frequencies is shown for weight normalized EDL muscle harvested from HSA-MCM/Numb(^f/f)^) or HSA-MCM/Numb^(f/f^/NumbL^(f/f)^ mice at 14-days after starting injections of tamoxifen or vehicle. Statistical analysis was performed with repeated measure ANOVA followed by Sidak’s multiple comparison test. F = 28.29, DFn= 6, DFd= 22; ***p< 0.001, force X frequency interaction ***p< 0.001; (B) Maximum force generated during a single twitch is shown. Statistical analysis was performed with one-way ANOVA with Tukey’s post-hoc test (F=10.07, DFn=3, DFd=13). *p<0.05; **p<0.01. N=3-6

**Figure 2.**
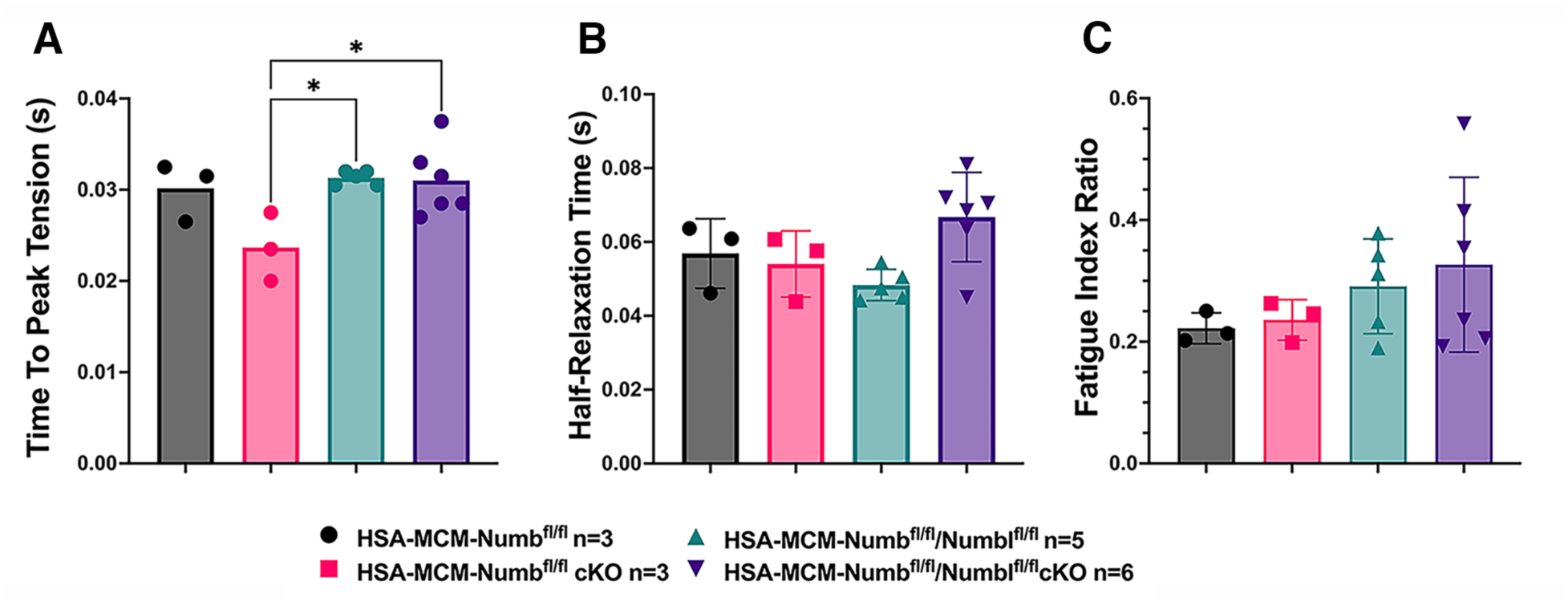
Time to peak tension (A), half-relaxation time (B), and fatigue index (C) are shown for HSA-MCM/Numb(^f/f)^) and double (HSA-MCM/Numb^(f/f^/NumbL^(f/f)^ mice) at 14 days after starting injections of tamoxifen or vehicle. Data were analyzed by one-way ANOVA with Tukey post-hoc test. (A) F=4.561, DFn=3, DFd=13; *p<0.05; (B) F=3.679, DFn=3,DFd=13 ; (C) F=0.9772, DFn=3, DFd=13. N=3-6.

To understand if relationships between frequency and tension were altered by Numb or Numb/NumbL cKO, data were re-plotted as percent of maximum tetanic tension on the Y axis and frequency on the X-axis. However, no shift in these curves was seen for either the Numb cKO or the Numb and NumbL cKO when compared to corresponding curves for vehicle-treated mice with normal expression levels of Numb and Numb (Supplementary Figure 3). These results confirm a critical role for Numb in force generating capacity of skeletal muscle.

### Analysis of Numb Splice Variants in the Myogenic Lineage

Given that Numb RNA can undergo alternative splicing to generate 4 different variants, and that alternative splicing shortens or removes protein-protein interaction domains, we sought to understand which splice variants of Numb were present in cells of the myogenic lineage in mice. Total RNA was extracted from mouse gastrocnemius muscle or from mouse C2C12 myoblasts then amplified using primers to sites flanking either exon 3 or exon 9 of mouse Numb mRNA (Supplementary Figure 1). No transcripts containing exon 9 were observed, and the majority of transcripts lacked exon 3, indicating that in skeletal muscle the primary form of Numb is that encoded by the shortest mRNA which lacks both exons 3 and 9. To gain some insight as to whether disease might alter Numb splice variants, this analysis was repeated for muscle at 7 days after sciatic nerve transection. No effect of denervation on splice variants present in muscle was observed (Supplementary Figure 1).

### Numb Immunoprecipitation

Having confirmed that Numb is responsible for most, if not all of the decrease in force production of muscle of our Numb/NumbL cKO mice, we sought to understand better how Numb participates in muscle force generating capacity. We determined the binding partners for Numb in cells of the myogenic lineage by proteomics analysis of protein complexes isolated by immunoprecipitation using anti-Numb antibodies. For these experiments, we used C2C12 myotubes which are multinucleated cell syncitia that express actin, myosin and acetyl choline receptors and can be induced to contract by electrical stimulation. Using the methods described above, we were able to precipitate almost quantitatively Numb proteins when detergent was included in the cell lysis buffer. Yields were low when detergent was removed which we infer indicates that in C2C12 myotubes, Numb is localized to membranes or is bound to membrane-bound proteins. Due to the incompatibility of LC/MS/MS analysis with most detergents commonly used in protein chemistry, choices of detergent that could be used were limited. After much optimization, we used a buffer containing a reduced amount of Triton-X100 (0.1%) for immunoprecipitation and washed the IP with TBS to remove as much Triton-X100 as possible. We also found that heating the samples at 37 °C for 1 h in 0.3% SDS was a mild yet effective elution method.

As shown in Supplemental Figure 4A, using this method, we were able to immunoprecipitate Numb from 5-day differentiated C2C12 myotubes. Numb IP was verified by Western blot using a rabbit monoclonal anti-Numb antibody whose specificity was previously validated by showing loss of Numb expression in C2C12 lysates treated with a specific vivo morpholino oligonucleotide [18]. Silver-stained SDS-PAGE gels revealed that immunoprecipitated proteins showed a uniform distribution of proteins across samples (Supplemental Figure 4B).

### Mass Spectrometry Data Analysis

Liquid chromatography (LC)/mass spectrometry (MS)/MS analysis of peptides generated by digestion of the immunoprecipitated samples by trypsin followed by database searches identified 17,394 peptides. Mass spectrometry raw data files have been deposited in an international public repository (MassIVE proteomics repository at https://massive.ucsd.edu/) under data set # MSV000089327. The raw data files may be accessed by ftp protocol at ftp://massive.ucsd.edu/MSV000089327/. Using a false discovery rate (FDR) of 1% for both peptide sequence and protein identification, 1122 proteins were identified. Among these, 437 proteins were removed from analysis since they were represented by a single peptide or had insufficient data (< 4 points /treatment) along with an additional 14 proteins that were derived from contaminants or added proteins. The analysis was conducted on 671 proteins that were represented by two or more peptides.

To identify the proteins that may interact with Numb among the 671 that passed the first screen, the following criteria were used: p<0.01, 2 or more peptides identified, and detection of the protein target in 4 or more of the samples analyzed. Using these criteria, 11 potential Numb binding proteins were identified (Table 1). A complete list of proteins for which peptide fragments were identified is given in Supplementary Table S1. Examples of mass spectra of representative peptides are shown in Supplementary Figures 5-10.

**Table 1.** Numb-interacting proteins identified by mass spectrometry.

| UniProt ID | Protein | Abundance ratio (Numb/control) | p value (control vs Numb) | Control count (out of 7) | Numb count (out of 7) |
| --- | --- | --- | --- | --- | --- |
| APCA4_MOUSE | Anaphase promoting subunit 4 | ∞ | 0 | 0 | 7 |
| NIPA_MOUSE | Nuclear-interacting partner of Alk | ∞ | 0 | 0 | 7 |
| PDE4D_MOUSE | cAMP specific 3-5-cyclic phosphodiesterase 4D | ∞ | 0 | 0 | 6 |
| WNK1_MOUSE | Serine-Threonine kinase Wnk1 | 289 | $5.4E^{-4}$ | 4 | 7 |
| SEPT7_MOUSE | Septin 7 | 82.9 | $3.6E^{-3}$ | 4 | 7 |
| APC 10_MOUSE | Anaphase promoting complex subunit 10 | ∞ | 0 | 0 | 5 |
| SEPT10_MOUSE | Septin 10 | ∞ | 0 | 0 | 4 |
| HOME3_MOUSE | Homer protein homologue | ∞ | 0 | 0 | 4 |
| TRAF7_MOUSE | E3 ubiquitin ligase Traf7 | ∞ | 0 | 0 | 4 |
| RUVB1_MOUSE | RuvB-like 1 | ∞ | 0 | 0 | 5 |
| SEPT2_MOUSE | Septin 2 | 379 | 0.01 | 5 | 7 |
| SEPT9_MOUSE | Septin 9 | 148 | 0.01 | 4 | 7 |

### GWAS and WGS Identify Relationships of Numb-binding Proteins and the Skeleton

We took two approaches to identifying potential links between these proteins and human physiology and disease. We first conducted searches of GWAS and WGS data (GWAS Catalog https://www.ebi.ac.uk/gwas/home; MSK-KP at https://msk.hugeamp.org/ [25]) for links between genetic variants in each of the genes encoding proteins identified as putative Numb-interacting proteins and diseases. Because of the extensive interactions of muscle and bone, our search included disorders of the skeleton and skeletal muscle. While no links to sarcopenia or other disorders of skeletal muscle were found, associations with several disorders of the skeleton were identified (Supplementary Table 2) that included: linkage between femoral neck bone mineral density (BMD) in women and phosphodiesterase 4D (PDE4D), and links between estimated BMD and Septin 9.

### Several Numb-Binding Partners are Implicated in Skeletal Muscle Function

A manually curated annotation was developed to understand the potential functional role of each of these proteins in skeletal muscle (Supplemental Table S2). Sources used were GeneCards [26] and literature searches using PubMed. These curated data were then compared to the phenotype reported for myotubes and myofibers depleted of Numb and NumbL [18] which includes reduced cell fusion, reduced mitochondrial function, delayed calcium transients and muscle weakness. Homer3 bound Numb in our IP/LC/MS/MS assay and is similar to Homer1, a protein which has been implicated in excitation-contraction coupling [27]. Septin 7 also bound Numb and was recently shown to cause multiple defects in skeletal muscle [28] that appear in many ways to phenocopy effects of KO of Numb and NumbL in myofibers [18], including altered mitochondrial morphology and muscle weakness. In total, we identified four proteins belonging to the septin family (Septin 2, 7, 9 and 10) as putative Numb-interacting proteins.

### Confirmation of Septin-7 Interaction with Numb

Septins are GTP-binding proteins that are involved in many cellular processes by functioning as scaffolds to recruit other proteins or to compartmentalize cellular domains [29]. Given the overlap in phenotype of skeletal muscle-restricted knockouts of Numb and Septin 7 [18, 28], we sought to confirm their association biochemically and spatially by confocal microscopy. Using differentiated C2C12 myotubes grown independently from those used for the LC/MS/MS analysis, we performed co-immunoprecipitation followed by western blot and confirmed the binding of Septin 7 to Numb (Figure 3 A). The pattern of immunostaining of Numb and Septin 7 was visualized by confocal microscopy (Figure 3 B-E). In longitudinal sections of TA muscle, Numb immunostaining was seen as intense wavy bands partially traversing the myofiber. Septin 7 immunostaining was seen as intense streaks along the length of the myofiber with fainter bands traversing the fiber. Panel 3 B in Figure 3 shows areas of Numb and Septin 7 co-localization.

**Figure 3.**
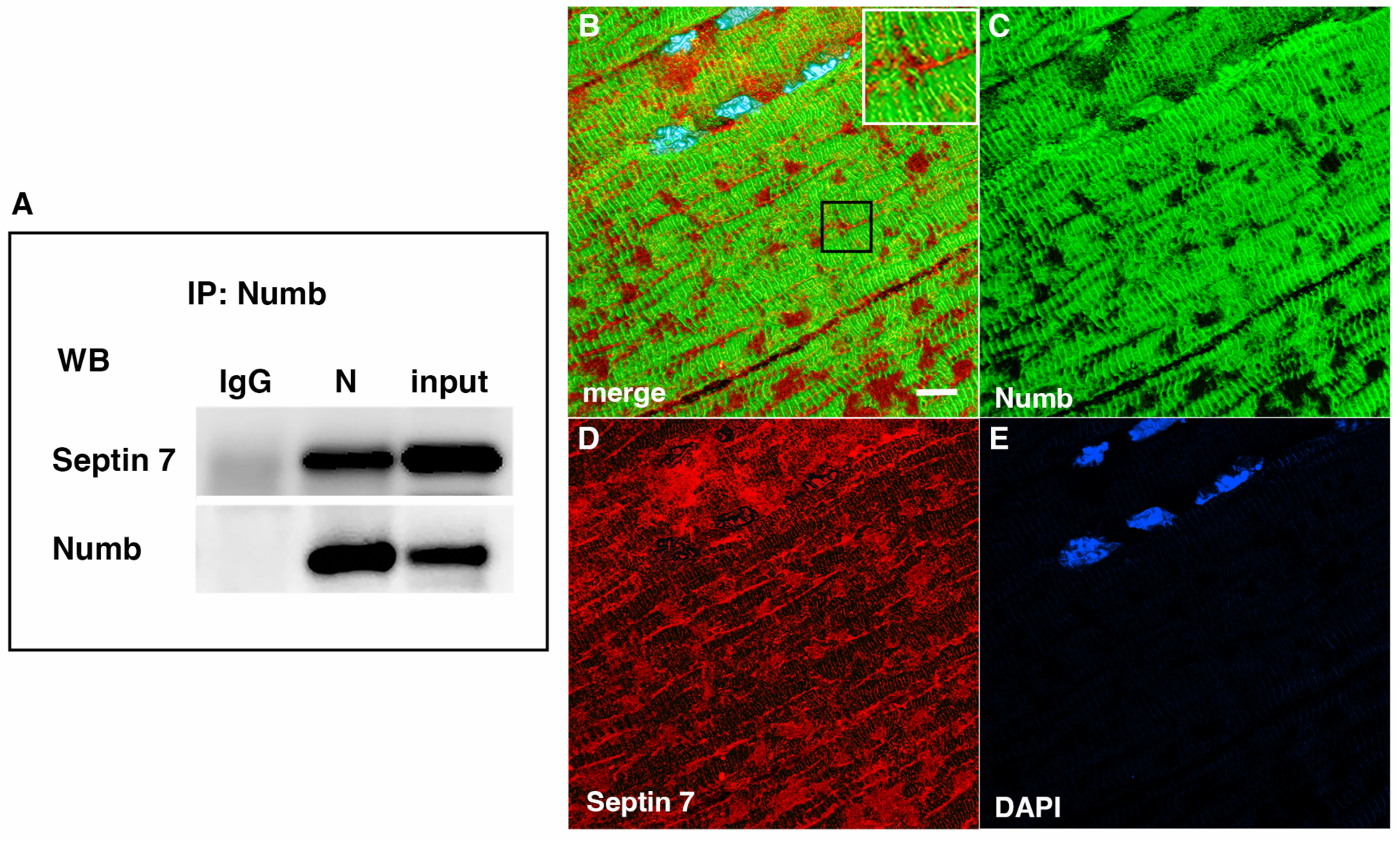
A Western blot showing a representative co-immunoprecipitation of Septin7 by anti-Numb antibody. IgG: immunoprecipitation with control Ig; N: immunoprecipitation with anti-Numb; input: original C2C12 lysate. The upper panel was probed with anti-Septin 7, the lower panel with anti-Numb. B-D. Immunolocalization of Numb and Septin 7 in muscle. Longitudinal cryosections of TA muscle from C57Bl6 mice were immunostained for Numb and Septin 7: (B) merged image; (C) anti-Numb antibody (green); (D) anti-Septin 7 antibody (red); (E) DAPI staining of nuclei (blue). Scale bar, 10 μm. The inset in A shows higher magnification (x2.3) of the area in the black square.

Myonuclei demonstrate specialization of gene expression profiles based on location within the myofiber. To understand if regional variations in Numb or Septin 7 expression might occur, we searched Myoatlas, a database of single nucleus sequencing data [30] (https://research.cchmc.org/myoatlas/). Numb was expressed in myonuclei throughout the myofiber (Figure 4 A) while, by contrast, Septin 7 expression was most abundant in myonuclei expressing acetylcholine receptor subunits (Figure 4 B). To understand if Septin 7 might be expressed at neuromuscular junctions, isolated myofibers were immunostained with anti-Septin 7 antibodies while acetylcholine receptors were labeled with α-bungarotoxin. Confocal microscopy imaging revealed greater intensity of Septin 7 immunolabeling at the neuromuscular junction (Figure 4 C).

**Figure 4.**
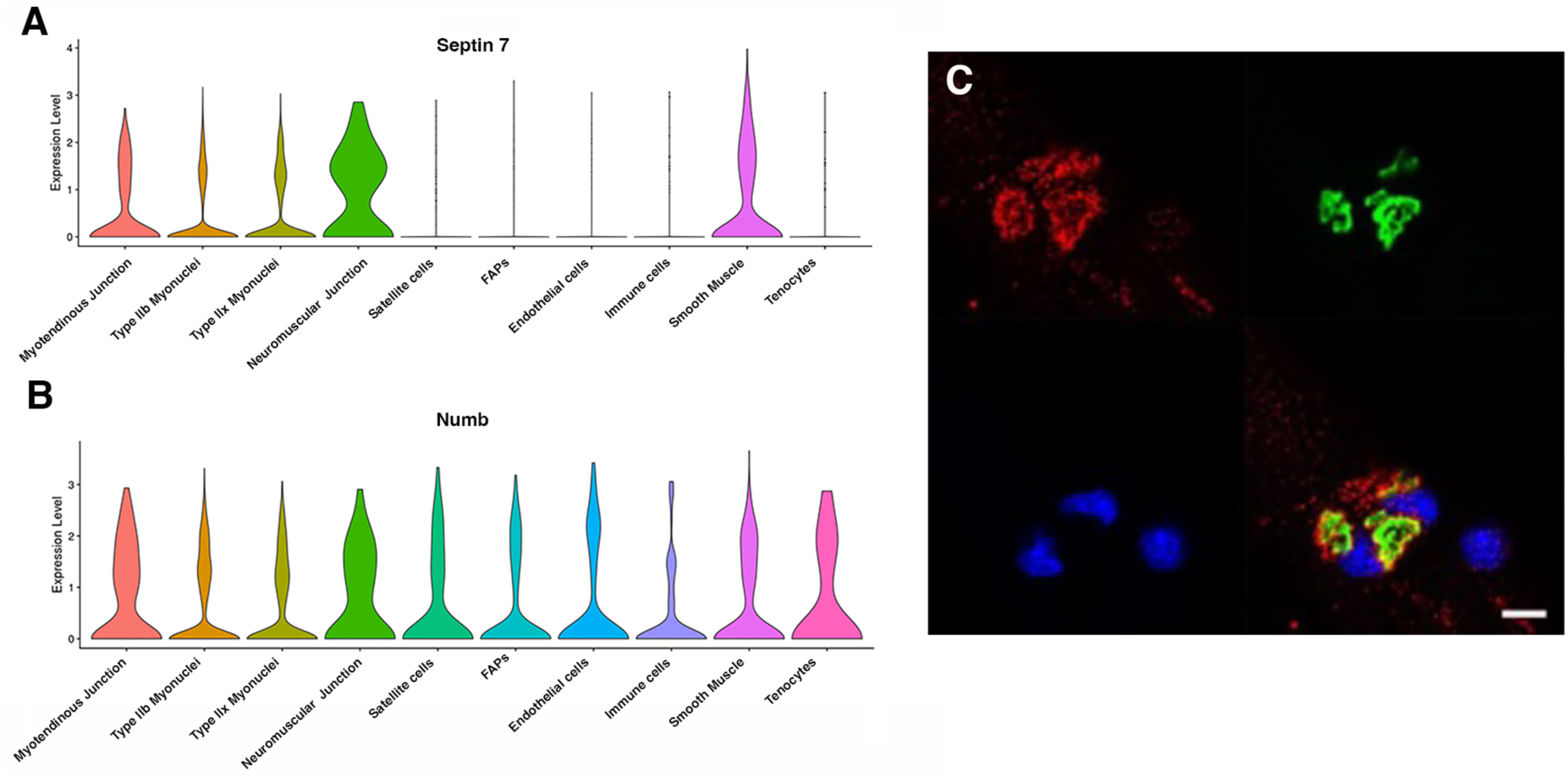
Septin 7 is enriched at neuromuscular junctions. A-B. Violin plots generated by Myoatlas showing abundance of Septin 7 (A) and Numb (B) mRNA in nuclei of TA muscle from 5-month old mice [30] C. Representative confocal microscopy image of the neuromuscular junction of an isolated myofiber immunostained with anti-Septin 7 (red) and with a fluorescently tagged α- bungarotoxin (green); nuclei were stained with DAPI (blue). Scale bar, 10 μm.

### Conditional Knockout of Numb and NumbL Perturbs Septin 7 Organization

The marked perturbation of sarcomeric ultrastructure observed in mice with conditional, inducible knockdown of Numb/NumbL in skeletal muscle led us to ask whether the highly ordered localization of Septin 7 was also lost when Numb levels were reduced. Localization of Septin 7 was determined by immunostaining followed by confocal microscopy using single myofibers isolated from mouse hindlimb muscle from HSA-MCM/Numb(f/f)/NumbL(f/f) mice at 14-days after induction of Numb/NumbL knockout with tamoxifen (Figure 5 D-F) or vehicle (Figure 5 A-C). In Numb/NumbL cKO myofibers, the pattern of immunostaining of Septin 7 was more punctate while with loss of the ordered, linear staining along the axis of fibers and traversing them (Figure 5 D-F).

**Figure 5.**
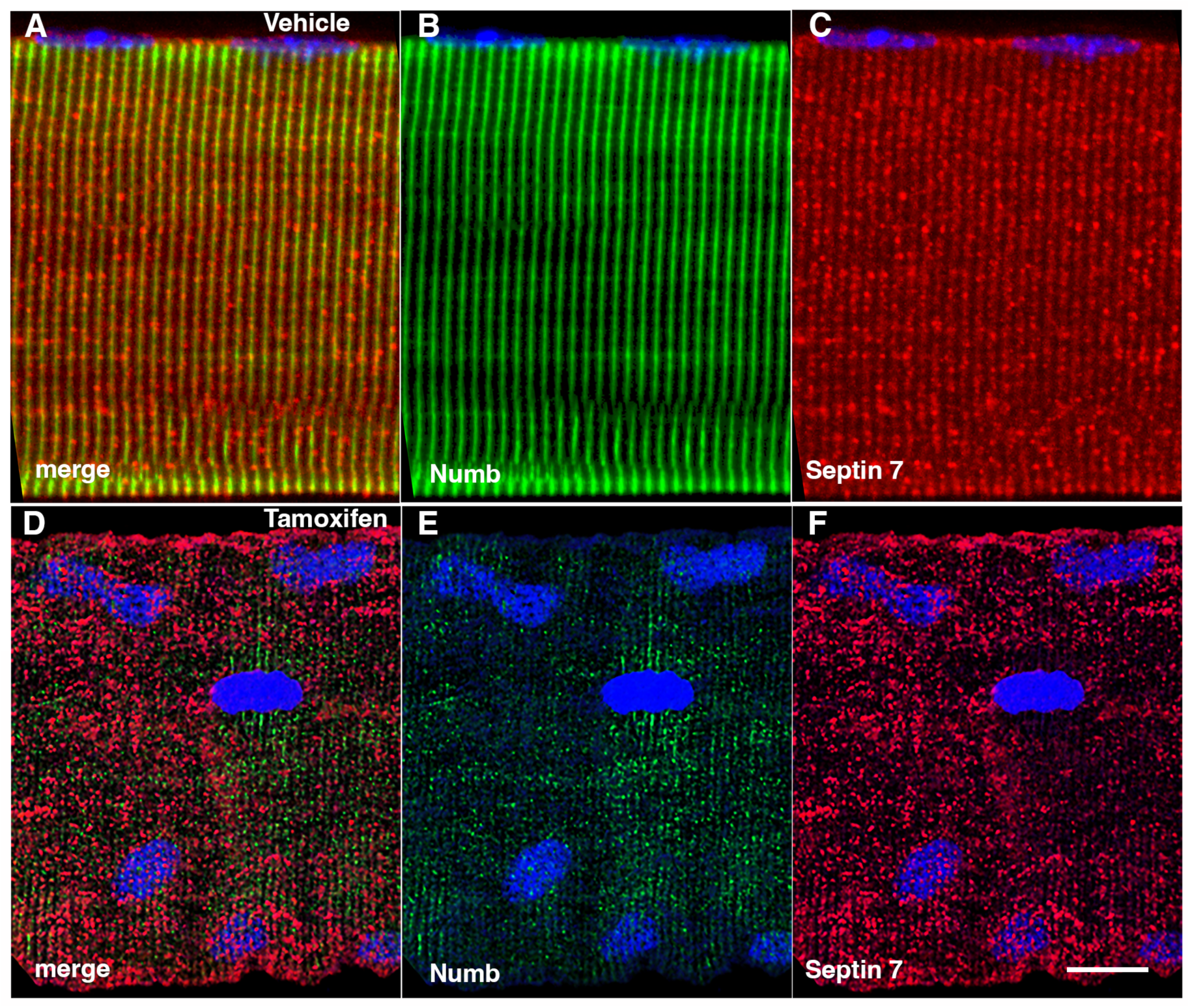
Representative confocal images of isolated mouse hindlimb fibers immunostained with anti-Numb (green) (B and E) and anti-Septin 7 antibodies (red) (C-F); merge: merged images (A and D). Nuclei were labeled with DAPI (blue). A-C: myofiber from a control mouse (vehicle treated); D-F, myofiber from a Numb cKO mouse (tamoxifen treated). Scale bar, 10 μm.

## DISCUSSION

The current study aimed to understand the molecular basis for the marked alterations in skeletal muscle ultrastructure, mitochondrial function, calcium release kinetics and force production caused by cKO of Numb and NumbL. Findings that a single, conditional and inducible KO of Numb in skeletal muscle fibers resulted in marked reductions of peak twitch and peak tetanic force that were similar, if not identical to those observed in a Numb/NumbL cKO confirm our conclusion that it was loss of Numb, rather than NumbL, that explained reduced muscle force production during *in-situ* physiologic testing in mice with a double Numb/NumbL KO [18]. The loss in force generating capacity of EDL was observed by *ex-*vivo physiological testing as soon as 14 days after the first injection of tamoxifen indicating that deterioration of muscle in response to depletion of Numb occurs rapidly. These findings do not, formally, test if NumbL contributes to homeostasis of adult skeletal muscle fibers although the very low-level expression of NumbL in adult skeletal muscle suggests that a role for this gene is unlikely [18].

Our approach to understanding how Numb contributed to the molecular physiology of skeletal muscle contractility relied on LC/MS/MS analysis of tryptic peptides of proteins immunoprecipitated from C2C12 myotubes using an anti-Numb antibody. By applying stringent criteria for identifying putative Numb binding partners, a total of 11 high-probability Numb binding proteins representing at least 6 different proteins or protein complexes was identified. None of these proteins were listed among the 209 known Numb-interacting proteins listed on the NCBI webpage for human Numb (https://www.ncbi.nlm.nih.gov/gene/8650#interactions). We believe these to be new, high-probability interactions, some of which may have muscle-specific roles.

The results support that conclusion that Septin 7 is an authentic binding partner of Numb within skeletal muscle fibers based on evidence that Numb pulls down Septin 7 from lysates of C2C12 myotubes, that Numb and Septin 7 colocalize within skeletal muscle fibers and that cKO of Numb and NumbL perturbs the distribution of Septin 7 immunostaining. The identification of three other septins as Numb binding partners is consistent with findings that septins form hetero-oligomers that self-organize into fibrils that can polymerize into sheets, filaments or rings [29]. The colocalization of Numb with Septin 7 is constrained to specific regions of the myofiber suggesting overlapping but distinct functions of each protein in organization of the sarcomere. For example, Septin 7 was detected without Numb immunostaining in several locations, including around the nuclear envelope, and in longitudinal streaks that traverse several sarcomeres. Why these proteins interact at some locations but not others is unclear. Possibilities are that splice variants of Septin 7 vary in their distribution based on proteins they interact with and that only some splice variants bind Numb. It may also be that interaction of Numb and Septin 7 is through a third yet to be identified protein that is localized near the triad, our proposed localization for Numb [18]. Further study is needed to determine the explanation for the distribution of these two critical proteins.

Our findings suggest that Numb may also interact with other septins such as septins 2, 9 and 10, which were also identified with a high level of confidence as Numb interacting proteins by our LC/MS/MS analysis. Our data do not allow us to determine if Numb binds directly to these septins. Septins contain highly conserved regions, and, consequently, if one such region of septin 7 interacts with Numb, then many septins would be expected to directly bind Numb through the same domain. However, because septins self-oligomerize, is possible that when Numb binds to one septin, antibodies against Numb could also pull down other septins present in the septin oligomer to which Numb is bound regardless of whether or not they are also bound by Numb.

Roles in muscle physiology of Homer3, RUVBL1, Wnk1 and TRAF7 remain uncertain. Homer1, a protein closely related to Homer3, has been implicated in interactions between the Cav1.2 subunit of L-type calcium channels and ryanodine receptors [27] and is expressed in skeletal muscle. Its expression levels also correlate with serum alkaline phosphatase levels (Supplemental Table 2 and [31]). TRAF7 is an E3 ubiquitin ligase regulated by MyoD that ubiquitinates NF-kB targeting it for proteasomal degradation [32] to facilitate myogenic differentiation. Zc3hc1is another E3 ubiquitin ligase highly expressed in skeletal muscle known for its ability to target Cyclin B for proteolysis by the ubiquitin-proteasome pathway.

Our interest in the role(s) of Numb in skeletal myofibers was stimulated in part by findings that its expression at the level of mRNA was reduced in older individuals [2], a finding we confirmed at the protein level in muscles from C57B6 mice [18]. The disorganization of Septin 7 immunostaining observed by day 14 after inducing Numb KD suggests that aging-related reduction of Numb expression may perturb organization of Septin 7 in a similar way though this prediction must be tested experimentally.

In conclusion, Numb binds Septin 7, a ubiquitously expressed protein involved in forming septin-based filaments, sheets and cages that serve as the ‘fourth component of the cytoskeleton’. The localization of these two proteins near the triad, together with binding of Homer 3 to Numb, strongly suggests that Numb and Septin 7 participate in organizing the triad though specific molecular interactions whose nature remains unclear. The interaction of Numb with septins have direct implications for understanding molecular physiology of skeletal muscle and broader implications for understanding roles of Numb and septins in biology.

