## Supplementary material for "Septin 7 Interacts With Numb To Preserve Sarcomere Structural Organization And Muscle Contractile Function": Suppl Material

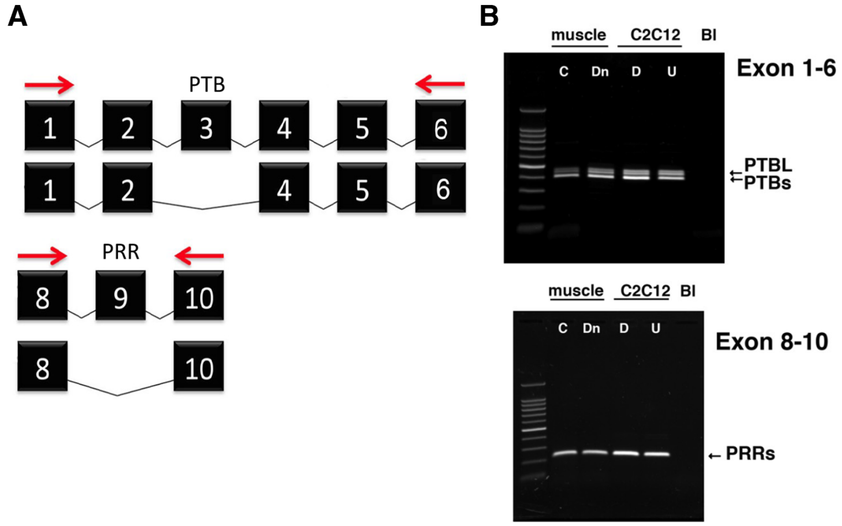


**Supplementary Figure 1.** Variants of Numb. (A) The PCR strategy used to detect Numb splice variants is shown. Red arrows depict the locations of the forward and reverse primers used. Exons (black boxes) are numbered 1-10. b. cDNA was prepared using total RNA isolated from denervated (Dn) or control (C) gastrocnemius muscle and from undifferentiated (U) and differentiated (D) C2C12 myoblasts and amplified using primers spanning exons 1-6 (upper panel) or 8-10 (lower panel). PCR products were analyzed by agarose gel electrophoresis. Gastrocnemius muscles were removed at 7 days after left sciatic nerve transection. Differentiated C2C12 cells were harvested at 5 days after transferring cells to differentiation media. PTBL and PTBs refers to phosphotyrosine binding domain with or without exon 3; PRRs: proline rich region, s refers to the PRR without exon 9 which is the only form present in muscle and myogenic cells.

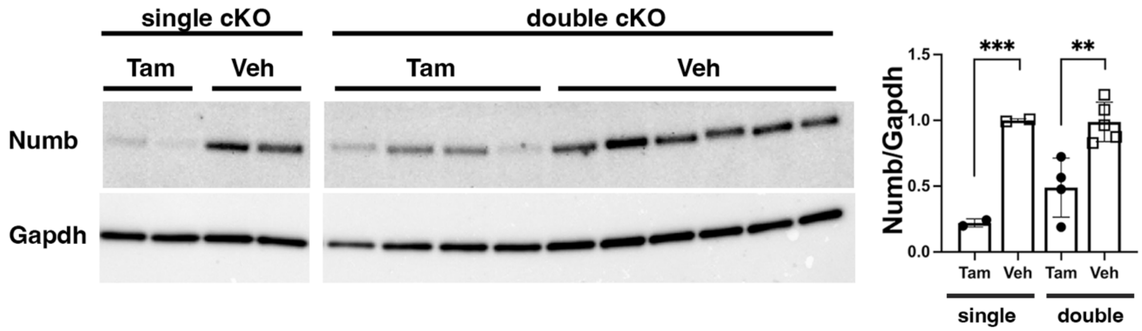


***Supplementary Figure 2.*** *Representative Western blot showing downregulation of Numb levels in EDL muscle from animals used for ex-vivo physiology experiments. Data were analyzed by unpaired t-test. **p<0.01. The lanes were cut from the same blot.*

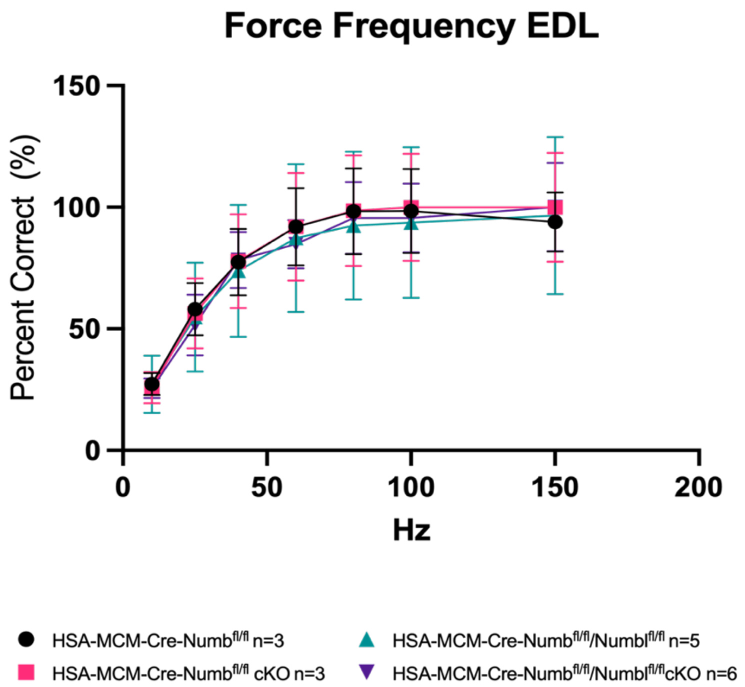


**Supplementary Figure 3.** Force-frequency data from Figure 1 were replotted for each group as percent maximal force produced for that group.

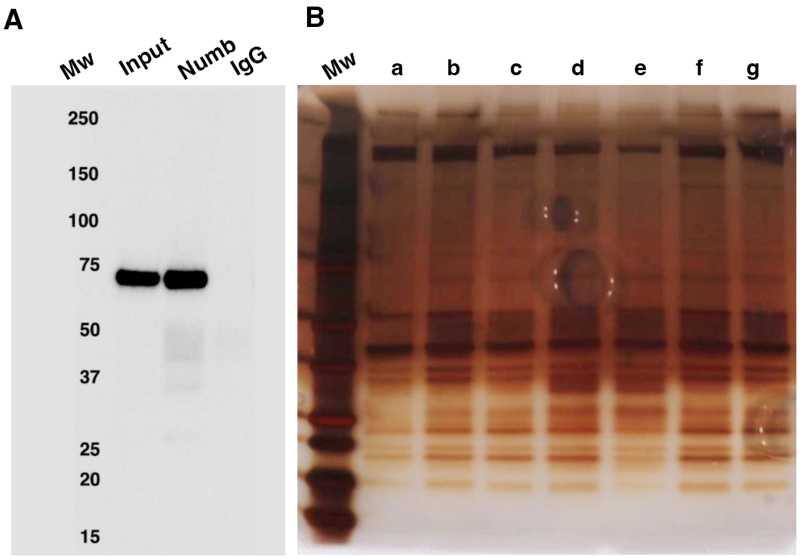


**Supplementary Figure 4.** Numb immunoprecipitation. (A) Representative Western blot of Numb immunoprecipitated from differentiated C2C12 myotubes. Input: original C2C12 lysate; Numb: goat anti-Numb IgG, control IgG: normal goat IgG. The blot was probed with a rabbit monoclonal anti-Numb antibody (B). Proteins recovered in each of the seven immunoprecipitations used for LC/MS/MS analysis were resolved by SDS-PAGE and visualized by silver staining.

### O55131|SEPT7_MOUSE Septin-7

#### Example Peptide Abundances for VNIIPLIAK (2+)

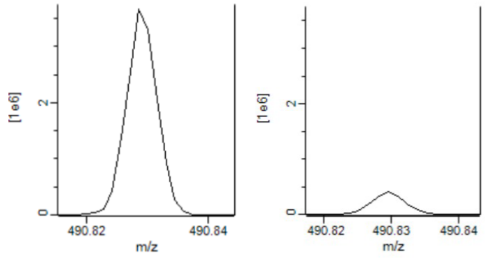


67 - Numb    67 - Control

| Peptides | Sequence coverage [%] | Protein Score | Abundance Ratio (Numb/ Control) | P-value (Control vs. Numb) |
|---|---|---|---|---|
| 32 | 67.7 | 210 | 82.9 | 3.6E-03 |

#### Example MS/MS Spectra

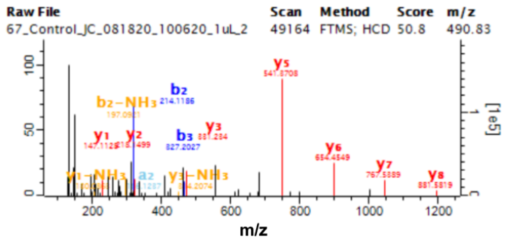


VNIIPLIAK (2+)

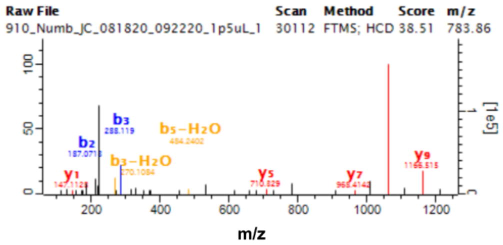


ADTLTPEECQQFK (2+)

**Supplementary Figure 5.** Mass spectra data for Septin 7 for representative IP sample (67) showing peptides abundance and mass spectra for representative peptides.

### Q01063|PDE4D_MOUSE cAMP-specific 3,5-cyclic phosphodiesterase 4D

#### Example Peptide Abundances for ELTHLSEMSR (3+)

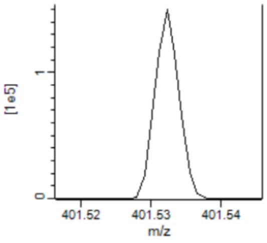


520 - Numb

No Signal

Control

| Peptides | Sequence coverage [%] | Protein Score | Abundance Ratio (Numb/ Control) | P-value (Control vs. Numb) |
|---|---|---|---|---|
| 5 | 9.6 | 14 | 1.0E+06 | 0 |

#### Example MS/MS Spectra

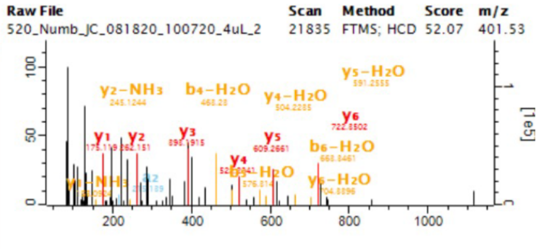


ELTHLSEMSR (3+)

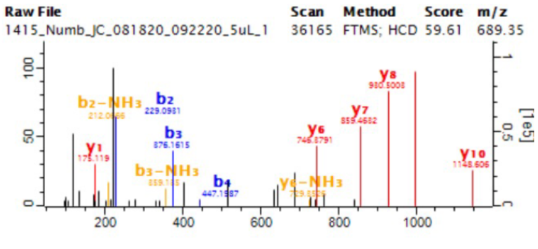


NNFAALTNLQDR (2+)

**Supplementary Figure 6**. Mass spectra data for PDE4E from representative IP sample (520) showing peptides abundance and mass spectra for representative peptides.

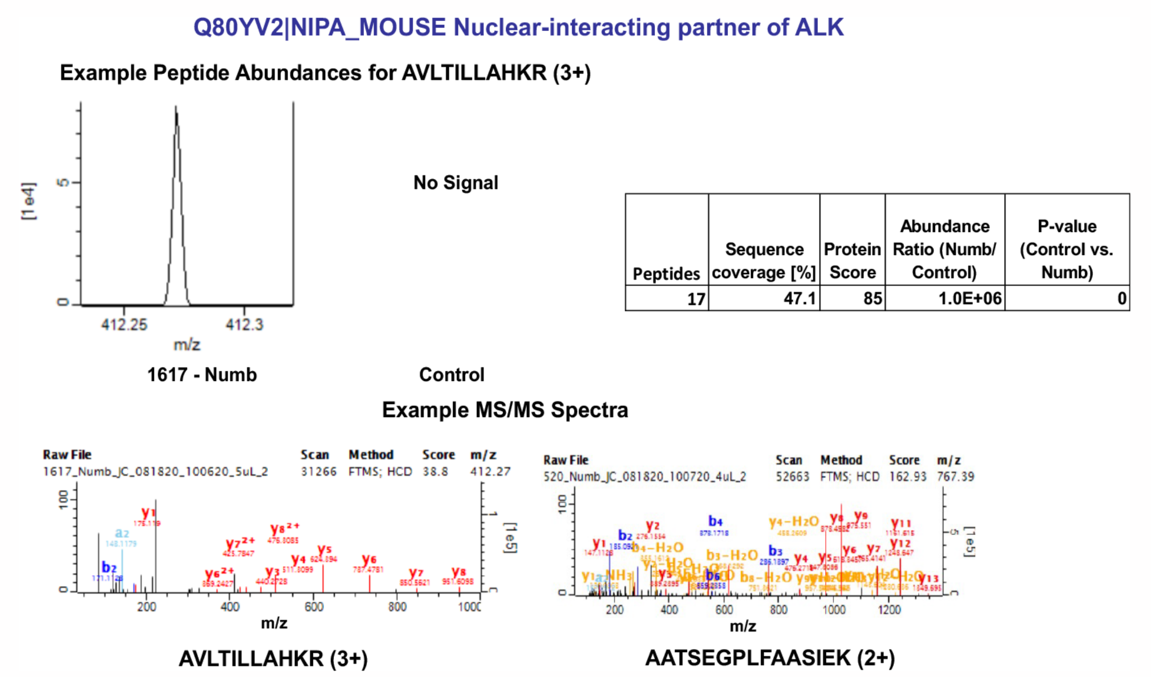


| Peptides | Sequence coverage [%] | Protein Score | Abundance Ratio (Numb/ Control) | P-value (Control vs. Numb) |
|---|---|---|---|---|
| 17 | 47.1 | 85 | 1.0E+06 | 0 |

**Supplementary Figure 7.** Mass spectra data for Nuclear interacting partner of ALK protein(NIPA) from representative IP sample (1616) showing peptides abundance and mass spectra for representative peptides.

### Q8BZQ7|ANC2_MOUSE Anaphase-promoting complex subunit 2

#### Example Peptide Abundances for IEELFSIIR (2+)

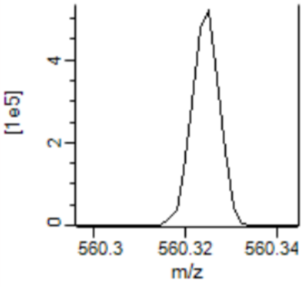


67 - Numb

No Signal

Control

| Peptides | Sequence coverage [%] | Protein Score | Abundance Ratio (Numb/ Control) | P-value (Control vs. Numb) |
|---|---|---|---|---|
| 27 | 36.2 | 142 | 1.0E+06 | 0 |

#### Example MS/MS Spectra

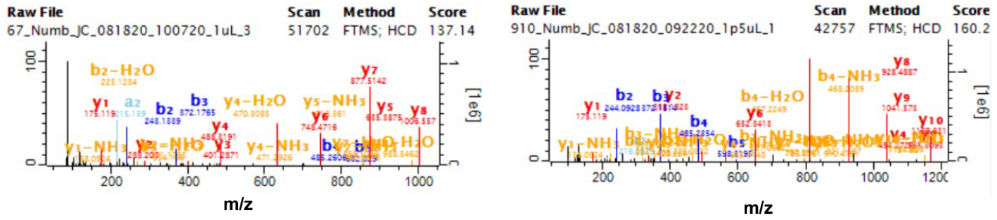


IEELFSIIR (2+)

DQQLIYSAGVYR (2+)

**Supplementary Figure 8.** Mass spectra data for ANC2 from representative IP sample (67) showing peptides abundance and mass spectra for representative peptides.

### Q91W96|APC4_MOUSE Anaphase-promoting complex subunit 4

**Example Peptide Abundances DLIALANTTGEVLLHR (3+)**

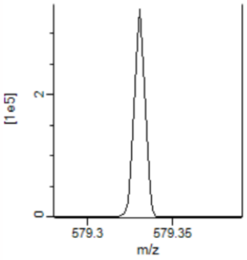


**No Signal**

| Peptides | Sequence coverage [%] | Protein Score | Abundance Ratio (Numb/ Control) | P-value (Control vs. Numb) |
|---|---|---|---|---|
| 26 | 38.5 | 97 | 1.0E+06 | 0 |

**1617 - Numb** **Control**

**Example MS/MS Spectra**

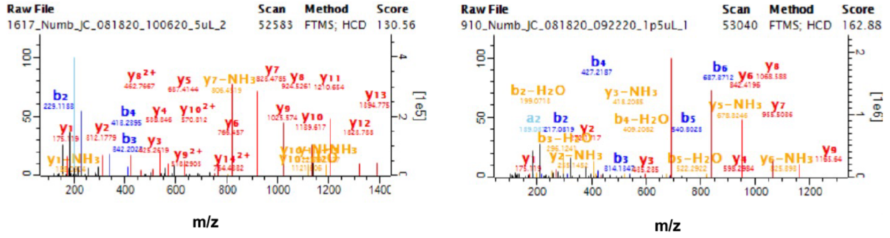


**DLIALANTTGEVLLHR (3+)** **ESPLLFPYYPR (2+)**

**Supplementary Figure 9.** Mass spectra data for APC4 from representative IP sample (67) showing peptides abundance and mass spectra for representative peptides.

**Q8K2H6|APC10_MOUSE Anaphase-promoting complex subunit 10**

**Example Peptide Abundances for VGNNFHNLQEIR (3+)**

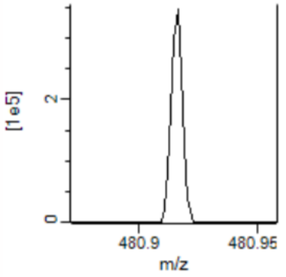


**520 - Numb**

**No Signal**

**Control**

| Peptides | Sequence coverage [%] | Protein Score | Abundance Ratio (Numb/ Control) | P-value (Control vs. Numb) |
|---|---|---|---|---|
| 4 | 31.4 | 11 | 1.0E+06 | 0 |

**Example MS/MS Spectra**

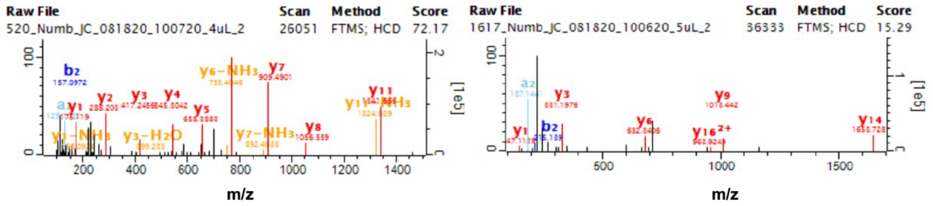


**Supplementary Figure 10.** Mass spectra data for APC10 from representative IP sample (67) showing peptides abundance and mass spectra for representative peptides.
